# Steroid hormone-dependent glial-neuronal interaction promotes brain development during *Drosophila* metamorphosis

**DOI:** 10.64898/2025.12.30.696973

**Authors:** Eisuke Imura, Naoki Okamoto, Naoki Yamanaka

## Abstract

Steroid hormones regulate various aspects of brain development in metazoans. In the fruit fly *Drosophila melanogaster*, the primary steroid hormone ecdysone enters the central nervous system (CNS) via Ecdysone Importer (EcI) in the blood-brain barrier (BBB) and induces brain development during metamorphosis. However, our understanding of the exact cell types that require ecdysone during CNS transformation is still limited. Here, we report that ecdysone-dependent glial-neuronal interaction promotes brain development during *Drosophila* metamorphosis. Unexpectedly, disrupting ecdysone signaling in glial cells caused more severe defects in CNS transformation than in neurons, suggesting an essential role of glia in ecdysone-dependent brain development. When *ecdysone receptor* was knocked down in mushroom body (MB) neurons, pruning of their larval-specific axonal lobes was disrupted as reported previously. In contrast, knockdown of *EcI* in the MB did not induce any discernible deficiency in neuronal remodeling, suggesting dispensability of EcI-mediated ecdysone entry into the MB. Consistent with this, the neuronal remodeling defects induced by *EcI* knockdown in the BBB were rescued by glial cell-specific overexpression of a transforming growth factor-β ligand *myoglianin* and an engulfment receptor *draper*, both of which are upregulated in glial cells in an ecdysone-dependent manner. Collectively, our findings suggest that ecdysone is primarily required in glial cells for CNS transformation during metamorphosis, elucidating a hormone-dependent glial-neuronal interaction that drives brain development.

## Introduction

The insect central nervous system (CNS) is transformed during metamorphosis, which is initiated and controlled by the steroid hormone ecdysone (1–4). Ecdysone signaling has been shown to regulate multiple aspects of CNS transformation, such as neuronal remodeling, differentiation, and programmed cell death, primarily through studies targeting ecdysone receptor (EcR), the nuclear receptor for ecdysone (5–11). For example, it is well established that remodeling of the mushroom body (MB), the center for associative learning in insects (12, 13), requires both *EcR* and its heterodimeric partner *ultraspiracle* in a cell-autonomous manner during *Drosophila* metamorphosis (9, 10, 14, 15). Ecdysone signaling is thus central to CNS metamorphosis, driving the transformation of the nervous system to accommodate adult-specific physiology and behavior.

Recently, we identified Ecdysone Importer (EcI), a membrane transporter required for ecdysone incorporation across cellular membranes (16). Importantly, EcI is indispensable for transcellular transport of ecdysone across the blood-brain barrier (BBB) during *Drosophila* brain development, demonstrating the necessity of the steroid ligand for EcR during CNS transformation (17). On the other hand, exact cell types that require ecdysone for transforming the neuronal network during metamorphosis still remain unexplored.

In this study, we targeted both *EcR* and *EcI* in various cell types to revisit the necessity of ecdysone signaling in different cellular populations within the *Drosophila* CNS during metamorphosis. To our surprise, glial-specific disruption of ecdysone signaling led to more severe impairments in CNS transformation than neuronal disruption, underscoring the indispensable contribution of glia to ecdysone-dependent brain development. We identified two ecdysone-inducible genes, transforming growth factor-β (TGF-β) ligand *myoglianin* (*myo*) and an engulfment receptor *draper* (*drpr*), which can together rescue neuronal remodeling defects induced by *EcI* knockdown in the BBB. Our results thus reveal an ecdysone-dependent glial-neuronal interaction essential for CNS transformation during insect metamorphosis.

## Results

### Ecdysone is required in glial cells for brain development during metamorphosis

As a first step toward identifying cell types that require EcI and ecdysone during CNS metamorphosis (Figure 1A and 1B), we conducted knockdown of *EcI* in entire neurons using pan-neuronal drivers (*Elav-GAL4* and *nSyb-GAL4*; Figure 1C and S1A). Despite the requirement of ecdysone entry into the CNS for brain development and pupation (17), knocking down *EcI* in entire neurons only showed minor CNS transformation defects in the pupal stage (Figure 1C), and most animals developed into either pharate adult or adult stages (Figure S1B). In contrast, knocking down *EcR* or overexpressing a dominant-negative form of *EcR* (*EcR-DN*) in entire neurons caused significant CNS transformation defects as visualized by the reduced brain lobe size, particularly when *nSyb-GAL4* was used, along with significant developmental lethality (Figure 1C and S1). Overall, these results are consistent with previous studies that showed necessity of *EcR* in different neuronal populations during *Drosophila* metamorphosis, whereas neuronal requirement of *EcI* remains less clear.

**Figure 1.**
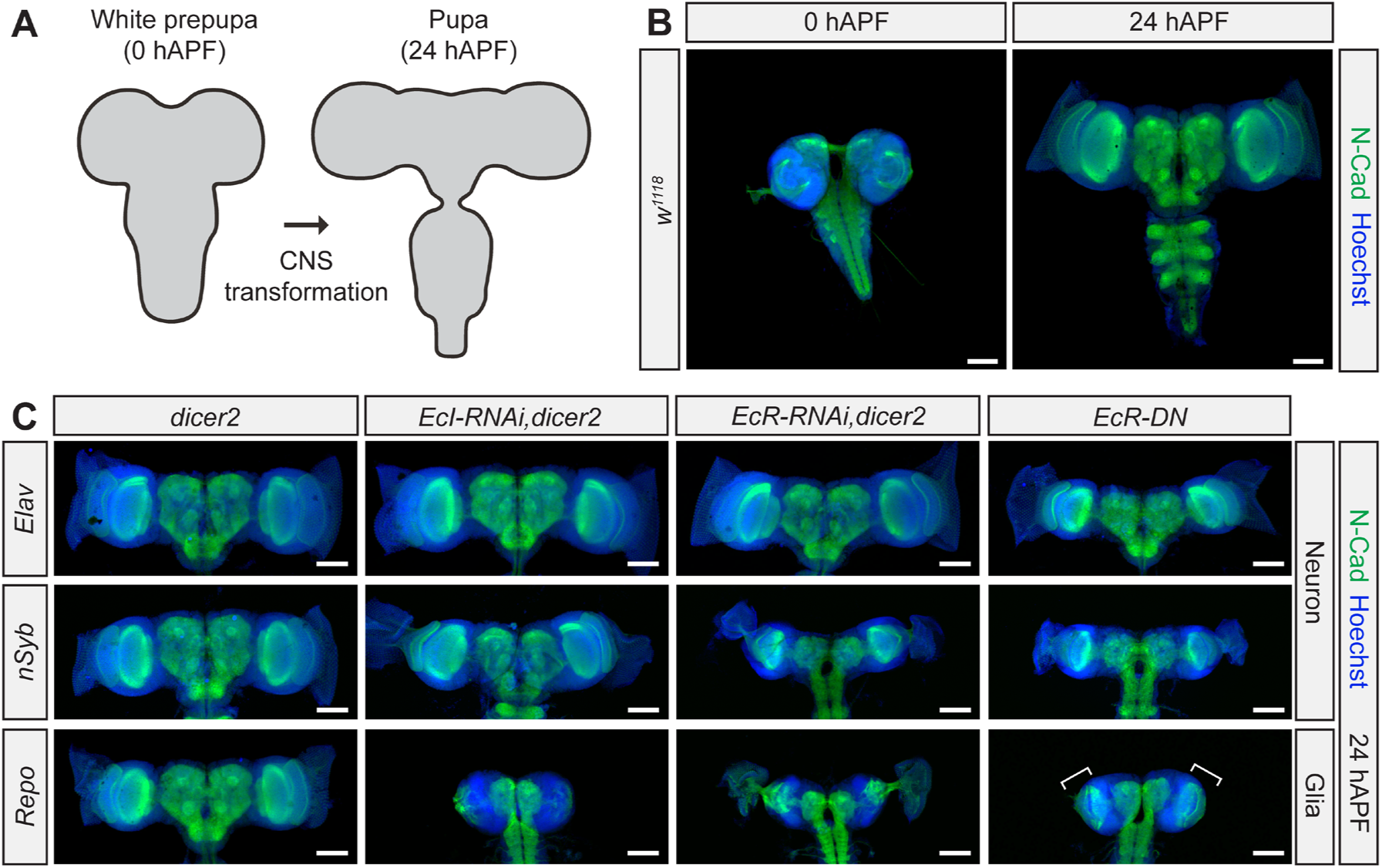
Ecdysone signaling in glial cells is required for CNS transformation during metamorphosis. (A) Schematic illustration of CNS morphology at the white prepupal stage (0 h after puparium formation or 0 hAPF) and pupal stage (24 hAPF). (B) CNS morphology at the white prepupal stage (0 hAPF) and pupal stage (24 hAPF) of the control strain *w^1118^* stained with anti-N-cadherin (N-Cad) antibody (green) and Hoechst 33342 (blue). Scale bars represent 100 μm. (C) Images of the pupal brain at 24 hAPF of control animals, as well as those expressing *EcI-RNAi*, *EcR-RNAi*, or a dominant-negative form of *EcR* (*EcR-DN*) in neurons or glial cells. *Elav-GAL4, UAS-dicer2* or *nSyb-GAL4, UAS-dicer2* and *Repo-GAL4, UAS-dicer2* were used to induce transgene expression in entire neurons and glial cells, respectively. Neuronal axons were stained with anti-N-Cad antibody (green), and nuclei were stained with Hoechst 33342 (blue). Differentiation of the optic lobe neuroepithelium in the brain of *Repo>EcR-DN* animals visualized by N-Cad staining is indicated by white brackets. Scale bars represent 100 μm.

We next knocked down *EcI* or suppressed *EcR* activities in glial cells. *EcI* knockdown in glial cells using *Repo-GAL4* blocked CNS transformation almost completely as expected, as it blocks ecdysone entry into the entire CNS through the BBB-forming glial cells (Figure 1C and S1A). To our surprise, both *EcR* knockdown and *EcR-DN* overexpression also caused CNS transformation defects when induced in glial cells, which were more severe than those caused by their induction in neurons (Figure 1C and S1A). The effect of suppressing EcR activity was strongest in *Repo>EcR-DN* animals, where the brain mostly retained the immature larval form 24 hours after puparium formation (hAPF), similar to *Repo>EcI RNAi* animals. The only major brain development observed in *Repo>EcR-DN* as compared to *Repo>EcI RNAi* animals was the differentiation of the optic lobe neuroepithelium (white brackets in Figure 1C). This is consistent with previous observations, as ecdysone has been shown to play important roles in timing cellular differentiation in the optic lobes (18–20). Overall, our results indicate that glial cells are the primary cell type that incorporate ecdysone through EcI to initiate brain development during *Drosophila* metamorphosis.

### Ecdysone receptor, but not Ecdysone Importer, is required in mushroom body neurons for remodeling

The MB is the center for associative learning in insects (12, 13). It is extensively remodeled during metamorphosis, and numerous studies have unequivocally shown that *EcR* is required for MB remodeling in a cell-autonomous manner during *Drosophila* metamorphosis (Figure 2A) (9, 10, 14, 15). Through *EcI* knockdown in the BBB-forming glial cells, we have previously shown that ecdysone entry into the CNS is also required for MB remodeling (17). Interestingly, we noticed that MB remodeling is disrupted not only by knocking down *EcI* but also by blocking EcR activity in glial cells: neuropile staining using anti-Fasciclin-II (Fas-II) antibody revealed that the larval-specific γ lobes of the MB were pruned and α/β neurons extended their axons at 24 hAPF in control animals, whereas this remodeling was blocked by knocking down *EcI* or blocking EcR activity in glial cells using *Repo-GAL4* (Figure 2B). This makes it unclear whether ecdysone is required as the EcR ligand in MB neurons or glia for promoting MB remodeling. We therefore first confirmed that EcR is highly expressed in Kenyon cells (KCs), the intrinsic MB neurons whose axons form the MB lobes, as previously reported (Figure 2C). Despite this abundant EcR expression, the ecdysone response element (EcRE)-driven LacZ reporter (*EcRE-LacZ*) (21) failed to detect any ligand-dependent EcR activity in KCs at pupariation (0 hAPF) or pupation (12 hAPF), when the ecdysone titer surges in the hemolymph (Figure 2D). We next knocked down *EcR* or *EcI* specifically in MB neurons using *OK107-GAL4* and visualized the MB by anti-Fas-II staining. In control animals, the larval-specific γ lobes were completely pruned and α/β lobes were extended at 24 hAPF, whereas this remodeling was blocked and the γ lobes remained intact in *EcR RNAi* animals, as reported previously (Figure 2E) (15, 22, 23). In contrast, *EcI RNAi* animals showed complete pruning of the MB γ lobes and axon extension of α/β neurons at 24 hAPF with no discernible defects (Figure 2E), suggesting that EcI-mediated ecdysone incorporation is not required in MB neurons for their remodeling. Furthermore, actively eliminating ecdysone out of MB neurons by overexpressing *Early gene at 23 (E23)*, an ATP-binding cassette transporter that has been suggested to export ecdysone out of cells (24–26), also failed to block MB remodeling (Figure 2E). To exclude the possibility that the observed phenotype reflects developmental delay rather than remodeling defects, MB morphology was also analyzed in adult flies of the same genotype three days after eclosion. The adult-specific γ-lobe morphology, characterized by medially extended axons, was consistently observed across all genotypes except those with *EcR* knockdown, in which the larval-specific vertical γ lobes persisted (Figure 2E). This observation further supports the essential role of *EcR* but not *EcI* in MB remodeling. The dispensability of ecdysone uptake in MB neurons was further confirmed by knocking down *EcI* in the MB of animals lacking three additional ecdysone importers, namely *EcI-2*, *-3*, and *-4* (Figure 2F) (27). Collectively, these results indicate that EcR activity, but not EcI-mediated ecdysone uptake, is essential for MB neurons to remodel during metamorphosis.

**Figure 2.**
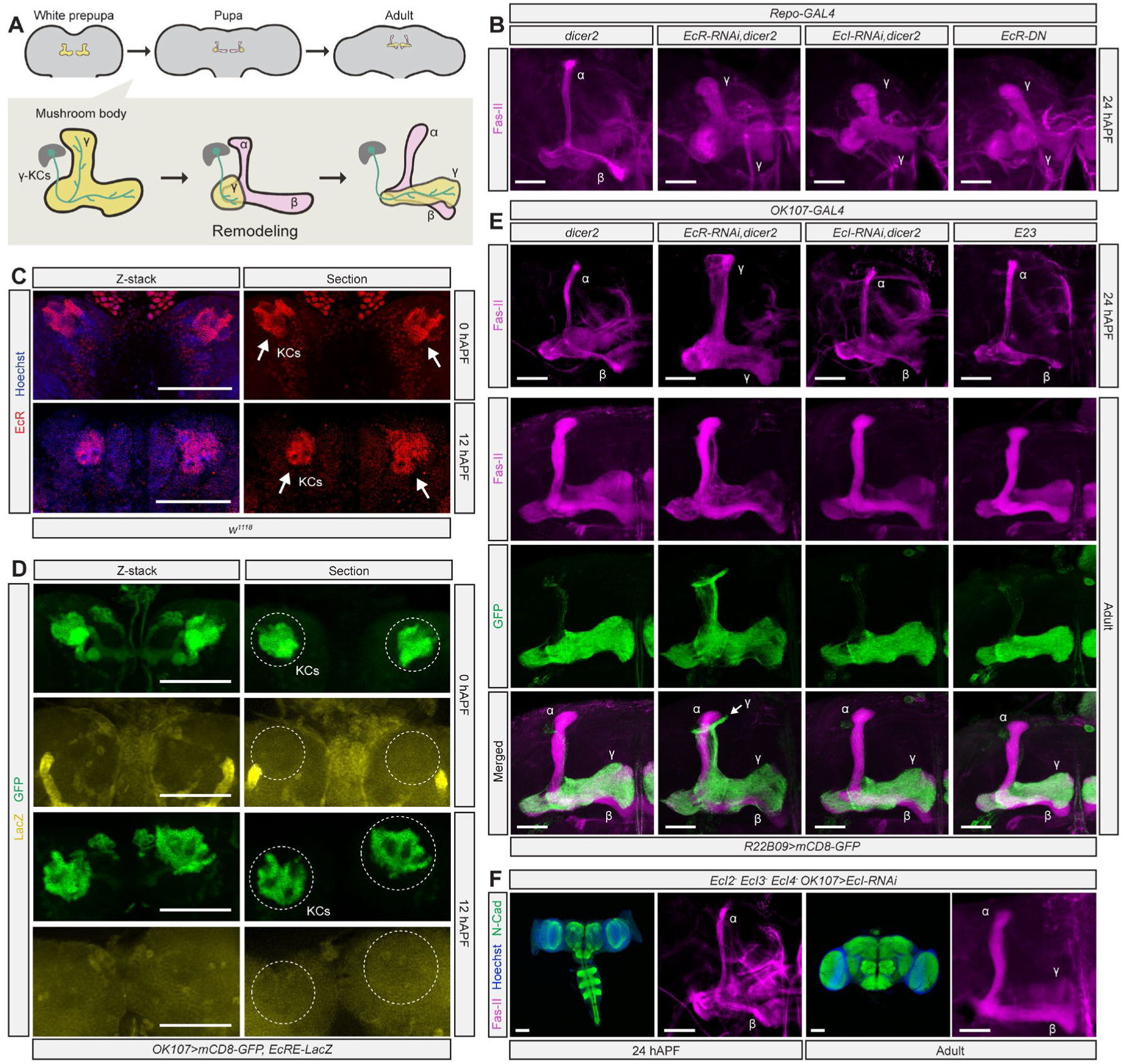
EcR, but not EcI, is required in the mushroom body for remodeling. (A) Schematic illustration of mushroom body (MB) remodeling during metamorphosis. α and β indicate the newly extending axonal lobes of a subset of Kenyon cells (KCs) called α/β neurons, whereas γ indicates the larval-specific axonal lobe of KC γ neurons. Cell bodies of γ neurons (γ-KCs) are located in the dark grey area. (B) Images of the MB at 24 h after puparium formation (24 hAPF) of control animals, as well as those expressing *EcI-RNAi*, *EcR-RNAi*, or a dominant-negative form of *EcR* (*EcR-DN*) in glial cells. *Repo-GAL4, UAS-dicer2* was used to induce transgene expression in glial cells. The MB lobes were visualized by anti-Fasciclin-II (Fas-II) antibody staining (magenta). Scale bars represent 25 μm. (C) EcR expression in the MB at 0 hAPF and 12 hAPF. EcR localization was visualized by anti-EcR antibody (red) in *w^1118^*. Nuclei were stained with Hoechst 33342 (blue). Allows indicate locations of KCs. Scale bars represent 100 μm. (D) Ligand-dependent EcR activity in the brain at 0 hAPF and 12 hAPF. The MB lobes were labeled by *OK107-GAL4*-driven expression of mCD8-GFP (green), whereas ligand-dependent EcR activity was visualized by the ecdysone response element-driven LacZ reporter (*EcRE-LacZ*; yellow). Dashed circles indicate locations of KCs. Scale bars represent 100 μm. (E) Images of the MB of pupae at 24 hAPF and adult males 3 days after eclosion. *OK107-GAL4* was used to induce *EcR-RNAi*, *EcI-RNAi*, or *E23* overexpression in the MB. The MB lobes were visualized by anti-Fas-II antibody staining (magenta). At the adult stage, the γ lobes were specifically labeled (green) by expressing *mCD8-GFP* under the control of the γ lobe-specific driver *R22B09-LexA*. Scale bars represent 25 μm. (F) Images of the pupal CNS and MB at 24 hAPF and adult males 3 days after eclosion. *OK107-GAL4* was used to induce *EcI-RNAi* in the MB of pupae in the *EcI2*, *EcI3*, and *EcI4* triple mutant background. Transheterozygotes of *EcI2*, *EcI3*, and *EcI4* null mutants and their deficiency strains were used. Neuronal axons were stained with anti-N-cadherin (N-Cad) antibody (green), nuclei were stained with Hoechst 33342 (blue), and the MB lobes were visualized by anti-Fas-II antibody staining (magenta). Scale bars represent 100 μm for the CNS images and 25 μm for the MB images.

The dispensability of EcI-mediated cellular ecdysone uptake in MB neuron remodeling is in clear contrast to cellular differentiation in the optic lobes, where an ecdysone pulse at the mid to late larval stage triggers differentiation of neuroepithelial cells into medulla neuroblasts (Figure S2A) (19, 20, 28, 29). Consistent with previous studies, *EcRE-LacZ* expression was clearly detected in the optic lobes, including the outer proliferation center (OPC) neuroepithelium labeled with *Ogre-GAL4* (Figure S2B) (17). Knocking down *EcI* or *EcR* using *Ogre-GAL4* suppressed this *LacZ* expression (Figure S2C and S2D) and reduced the size of the optic lobes (Figure S2E and S2F), which was also observed by OPC-specific overexpression of *E23* (Figure S2E and S2F). These results indicate that EcR requires its ligand ecdysone in the optic lobe neuroepithelium to induce cellular differentiation in a timely manner. Altogether, our results suggest that the ligand dependence of EcR activity varies among different cell types within the CNS, and EcI-mediated ecdysone incorporation is mainly required in glial cells rather than in KCs for MB remodeling during *Drosophila* metamorphosis.

### Ecdysone-dependent induction of the TGF-β ligand myoglianin in glial cells promotes CNS transformation

What are the factors that are induced by ecdysone in glial cells and promote neuronal remodeling? We first focused on *myo*, a TGF-β ligand-encoding gene expressed in glial cells and muscles during *Drosophila* development (30). It is known that glia-derived myo upregulates *EcR* expression in the MB and drives its remodeling following the late-larval ecdysone peak (Figure 3A) (31–33). Moreover, a previous study has suggested that *myo* expression in glial cells requires EcR activity (34), raising the possibility that myo is the factor that mediates glial-neuronal interaction induced by ecdysone.

**Figure 3.**
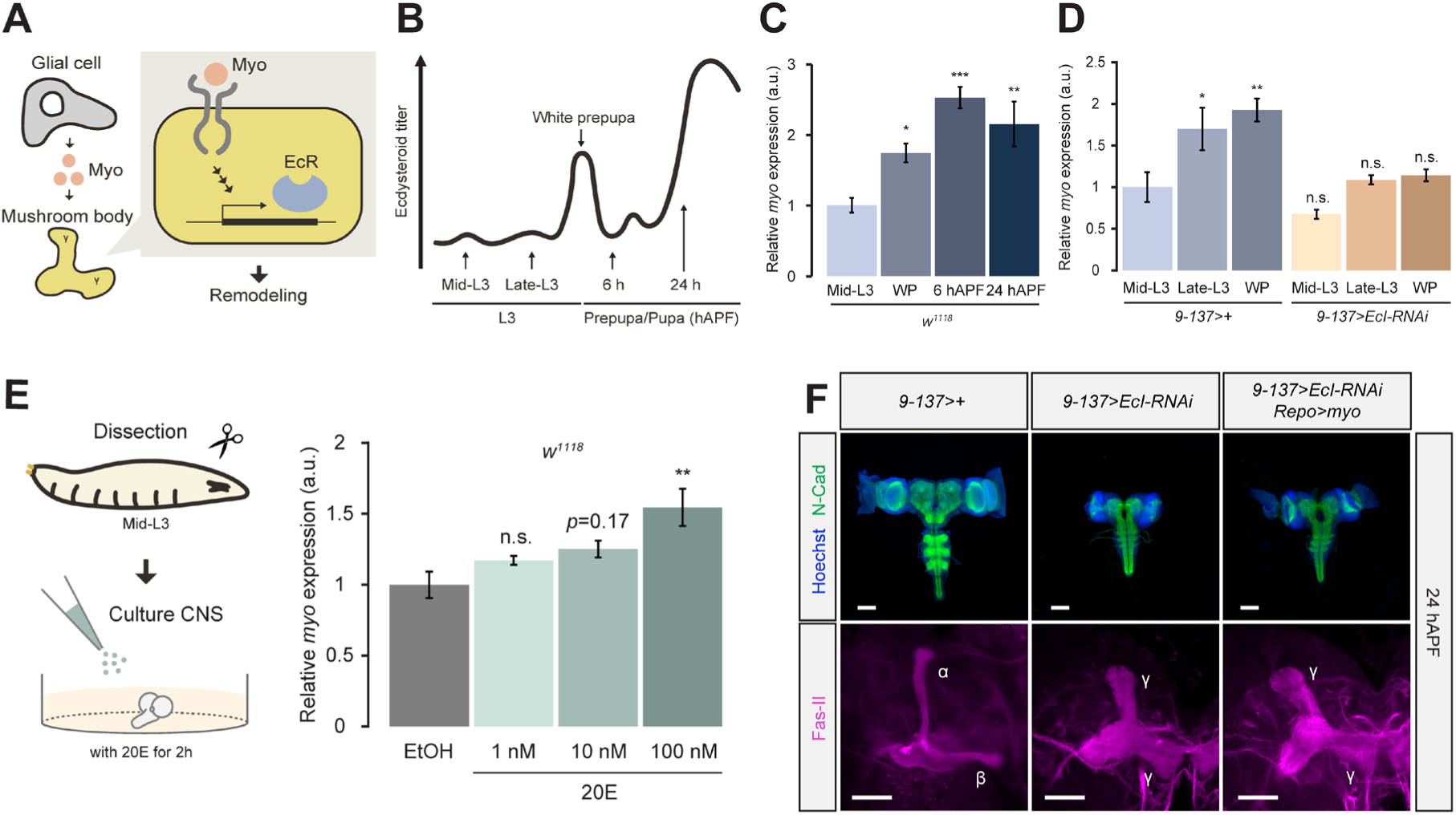
*Myoglianin* expression in glial cells is induced by ecdysone and promotes CNS transformation. (A) Schematic illustration of mushroom body (MB) remodeling induced by glia-derived myoglianin (myo). (B) Schematic diagram of fluctuations of the hemolymph ecdysone titer during early metamorphosis. hAPF, hours after puparium formation. (C, D) Relative expression levels of *myo* in the CNS during early metamorphosis, as assessed by qRT-PCR. The control strain *w^1118^* was used for C, whereas *9-137-LexA* was crossed to either *w^1118^*or *LexAop-EcI-RNAi* in D. Values are relative to that of *w^1118^*(C) or *9-137>+* (D) at the mid-L3 stage. a.u., arbitrary unit; WP, white prepupa. (E) Relative expression levels of *myo* in the CNS of mid-L3 *w^1118^* larvae cultured *ex vivo* for 2 h with various concentrations of 20-hydroxyecdysone (20E). Values are relative to that of the solvent control (EtOH). (F) Images of the pupal CNS and MB at 24 h after puparium formation (24 hAPF). *9-137-LexA* was used to induce *EcI-RNAi* in the BBB, whereas *Repo-GAL4* was used to overexpress *myo* in glial cells. Neuronal axons were stained with anti-N-cadherin (N-Cad) antibody (green), nuclei were stained with Hoechst 33342 (blue), and the MB lobes were visualized by anti-Fasciclin-II (Fas-II) antibody staining (magenta). Scale bars represent 100 μm for top and 25 μm for bottom panels. One-way ANOVA with post-hoc Dunnett’s test was used for C and E. Two-way ANOVA with post-hoc Dunnett’s test was used for D. ∗*p* ≤ 0.05, ∗∗*p* ≤ 0.01, ∗∗∗*p* ≤ 0.001, and n.s. for non-significant (p > 0.05). All values are the means ± standard error (*n* = 3).

To confirm that *myo* expression in glial cells indeed requires ecdysone, we first investigated *myo* expression levels in the CNS during *Drosophila* development. Expression of *myo* in the CNS is upregulated at pupariation when the hemolymph ecdysone titer surges, and the expression level remains high thereafter (Figure 3B and 3C). This upregulated *myo* expression at pupariation was suppressed in animals where ecdysone entry into the CNS was blocked by BBB-specific *EcI* knockdown (Figure 3D), suggesting that the upregulation of *myo* expression in glial cells during metamorphosis requires ecdysone. Ecdysone-dependent induction of *myo* expression in glial cells was further tested by CNS culture experiments, where the CNS was dissected from mid-third instar larvae and incubated *ex vivo* in the presence of ecdysone at concentrations that can induce expression of other, well characterized ecdysone-inducible genes (Figure 3E and S3). Expression of *myo* was induced by ecdysone in a dose-dependent manner, further suggesting that the upregulation of *myo* expression in glial cells during *Drosophila* metamorphosis is ecdysone-dependent.

To test if myo production in glia is sufficient to induce CNS transformation, we overexpressed *myo* in glial cells while blocking ecdysone entry into the CNS by combining *9-137-LexA/LexAop-EcI RNAi* and *Repo-GAL4/UAS-myo*. Overexpression of *myo* in glial cells partially rescued CNS morphology and prepupal lethality, suggesting that myo indeed mediates ecdysone-dependent CNS transformation during *Drosophila* metamorphosis (Figure 3F and S4). Importantly, glial overexpression of *myo* also restored EcR-B1 isoform expression in KCs, consistent with the previously reported function of myo in inducing *EcR* expression in the MB (Figure S5) (32). Despite all these changes, however, *myo* overexpression in glial cells did not rescue the MB remodeling defects (Figure 3F). Overall, these results indicate the presence of additional ecdysone-inducible factor(s) in glial cells that promote neuronal remodeling.

### Myoglianin and the engulfment receptor Draper cooperatively mediate ecdysone-dependent glial activities to promote CNS transformation

Genetic response of *Drosophila* glial cells to ecdysone is not limited to *myo* expression (35). Upon further pursuit of glial factors responsible for ecdysone-dependent neuronal remodeling, we focused on a single-pass transmembrane receptor Draper (Drpr), whose expression in glial cells requires EcR activity (36, 37). It has been shown that pruning of MB γ axons, which involves glial engulfment and clearing of axonal debris, is dependent on *drpr* expression in glial cells (Figure 4A) (38, 39). Axonal chemokine-like protein presented extracellularly on MB γ axons likely acts as the ligand for Drpr, and this ligand-receptor interaction promotes glial phagocytosis of neurons (Figure 4A) (40–42). There are three isoforms of Drpr in *Drosophila*, and only Drpr isoform-I (Drpr-I) is known to be involved in glial engulfment of MB neurons (43). To determine whether *drpr-I* expression requires ecdysone, we first investigated its expression levels in the *Drosophila* CNS during development. Similar to *myo*, expression of *drpr-I* in the CNS is upregulated at pupariation, which was diminished in animals where ecdysone entry into the CNS was blocked (Figure 4B). Moreover, *drpr-I* expression was induced by ecdysone in a dose-dependent manner in the CNS culture experiment, further suggesting that the glial expression of *drpr-I* during *Drosophila* metamorphosis is ecdysone-dependent (Figure 4C).

**Figure 4.**
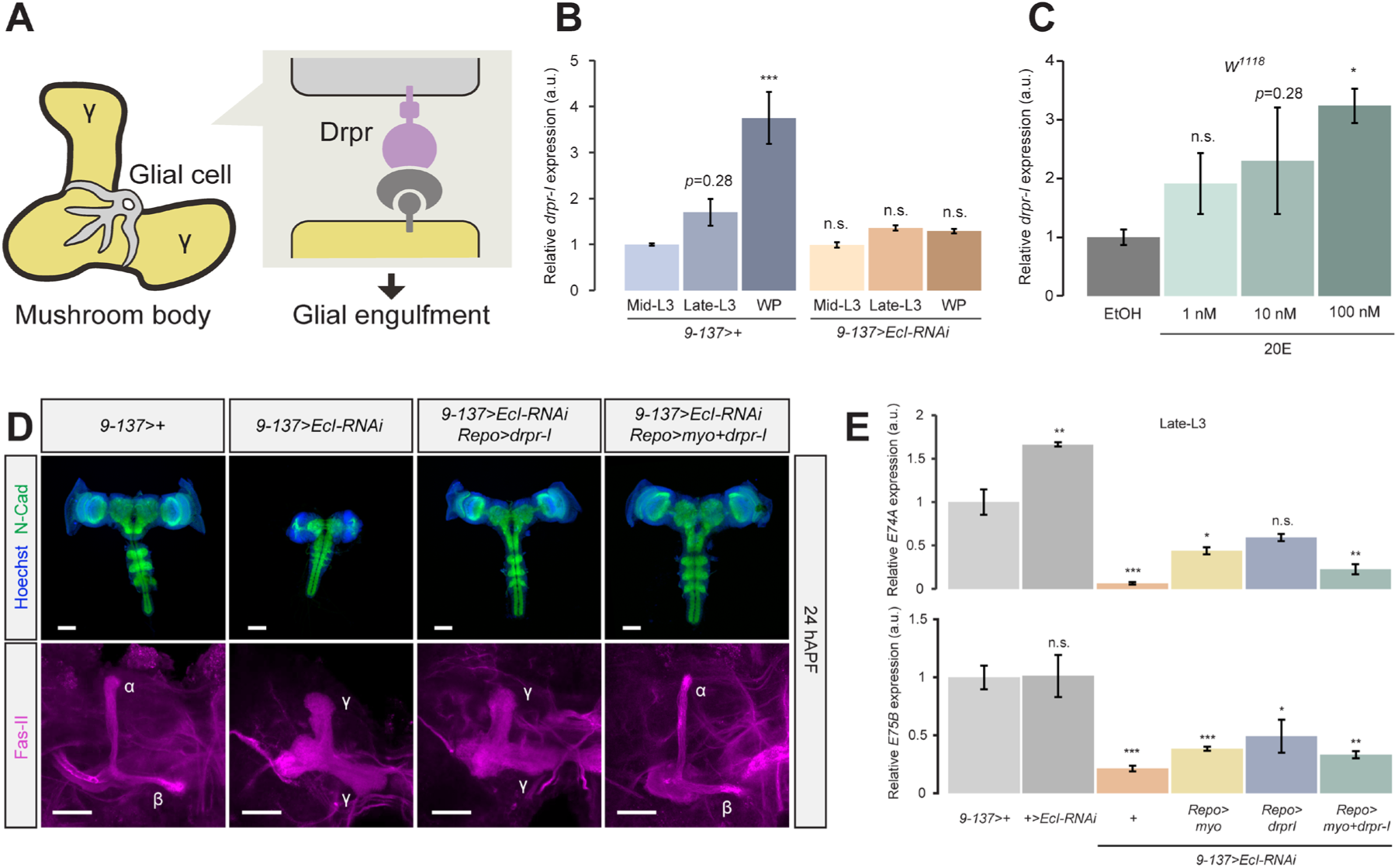
*Draper* expression in glial cells is induced by ecdysone and promotes CNS transformation in cooperation with *myoglianin*. (A) Schematic illustration of the Draper (Drpr) function in bridging mushroom body (MB) γ neurons and engulfing glial cells. (B) Relative expression levels of *draper isoform-I* (*drpr-I*) in the CNS during early metamorphosis, as assessed by qRT-PCR. *9-137-LexA* was crossed to either *w^1118^* or *LexAop-EcI-RNAi*. Values are relative to that of *9-137>+* at the mid-L3 stage. a.u., arbitrary unit; WP, white prepupa. (C) Relative expression levels of *drpr-I* in the CNS of mid-L3 *w^1118^* larvae cultured *ex vivo* for 2 h with various concentrations of 20-hydroxyecdysone (20E). Values are relative to that of the solvent control (EtOH). (D) Images of the pupal CNS and MB at 24 h after puparium formation (24 hAPF). *9-137-LexA* was used to induce *EcI-RNAi* in the BBB, whereas *Repo-GAL4* was used to overexpress *drpr-I* or *myoglianin* (*myo*)+*drpr-I* in glial cells. Neuronal axons were stained with anti-N-cadherin (N-Cad) antibody (green), nuclei were stained with Hoechst 33342 (blue), and the MB lobes were visualized by anti-Fasciclin-II (Fas-II) antibody staining (magenta). Scale bars represent 100 μm for top and 25 μm for bottom panels. (E) Relative expression levels of ecdysone-inducible genes (*E74A* and *E75B*) in the CNS at the late L3 stage (48 hours after L3 ecdysis), as assessed by qRT-PCR. *9-137-LexA* was used to induce *EcI-RNAi* in the BBB, whereas *Repo-GAL4* was used to overexpress *drpr-I* or *myo*+*drpr-I* in glial cells. Values are relative to that of *9-137>+*. Two-way ANOVA with post-hoc Dunnett’s test was used for B. One-way ANOVA with post-hoc Dunnett’s test was used for C and E. ∗*p* ≤ 0.05, ∗∗*p* ≤ 0.01, ∗∗∗*p* ≤ 0.001, and n.s. for non-significant (p > 0.05). All values are the means ± standard error (*n* = 3).

We next overexpressed *drpr-I* either alone or in combination with *myo* in glial cells while blocking ecdysone entry into the CNS and observed CNS morphology and MB remodeling. Co-overexpression of *myo* and *drpr-I* in glial cells drastically rescued CNS morphology, while *drpr-I* single overexpression only partially rescued the morphological defects (Figure 4D and S4A). Importantly, the MB remodeling defect caused by blocking ecdysone entry into the CNS was rescued by simultaneous overexpression of *myo* and *drpr-I*, but not by *drpr-I* single overexpression (Figure 4D). Likewise, prepupal lethality was dramatically rescued when both *myo* and *drpr-I* were overexpressed together (Figure S4B). Expression levels of ecdysone-inducible genes such as *E74A* and *E75B* in the CNS were suppressed in all animals with BBB-specific *EcI* knockdown (except a slight recovery in those carrying *Repo-GAL4/UAS-drpr-I*), confirming that inhibition of ecdysone entry into the CNS is not compromised by glial overexpression of transgenes (Figure 4E).

As *myo* overexpression in glial cells restored EcR-B1 expression in KCs (Figure S5), we reasoned that the MB remodeling defect caused by blocking ecdysone entry into the CNS may be rescued by combining glial overexpression of *drpr-I* and MB-specific overexpression of *EcR-B1*. Indeed, the MB remodeling defect caused by *EcI* knockdown in glial cells was rescued when *drpr-I* and *EcR-B1* were expressed simultaneously, but not separately, in glia and the MB, respectively (Figure S6). Altogether, our results indicate that ecdysone-inducible expression of *myo* and *drpr-I* in glial cells is the primary driver of neuronal remodeling and CNS transformation during *Drosophila* metamorphosis.

## Discussion

In the current study, we presented evidence suggesting that the insect steroid hormone ecdysone mainly acts on glial cells to promote CNS transformation during *Drosophila* metamorphosis. Although previous studies have clearly shown that EcR, the nuclear receptor for ecdysone, is required in neurons for their remodeling in a cell-autonomous manner (9, 10, 14, 15), necessity of its ligand ecdysone has never been thoroughly investigated, mainly due to the lack of tools for such studies. However, our recent identification of EcI, the membrane transporter required for ecdysone incorporation across cell membranes (16, 17), enabled us to examine requirements of cellular incorporation of ecdysone into distinct populations of cells within the *Drosophila* CNS. To our surprise, our results suggest that EcI-mediated ecdysone entry is not required in certain neuronal populations, such as those in the MB, for their remodeling during CNS transformation. Instead, ecdysone acts on glial cells to induce expression of two genes, a TGF-β ligand *myo* and an engulfment receptor *drpr*, which act together to induce MB remodeling and CNS transformation (Figure 5). Altogether, our results challenge the general assumption that ecdysone is universally required in neurons for their remodeling and instead propose that a hormone-dependent, glial-neuronal interaction primarily drives brain development during insect metamorphosis.

**Figure 5.**
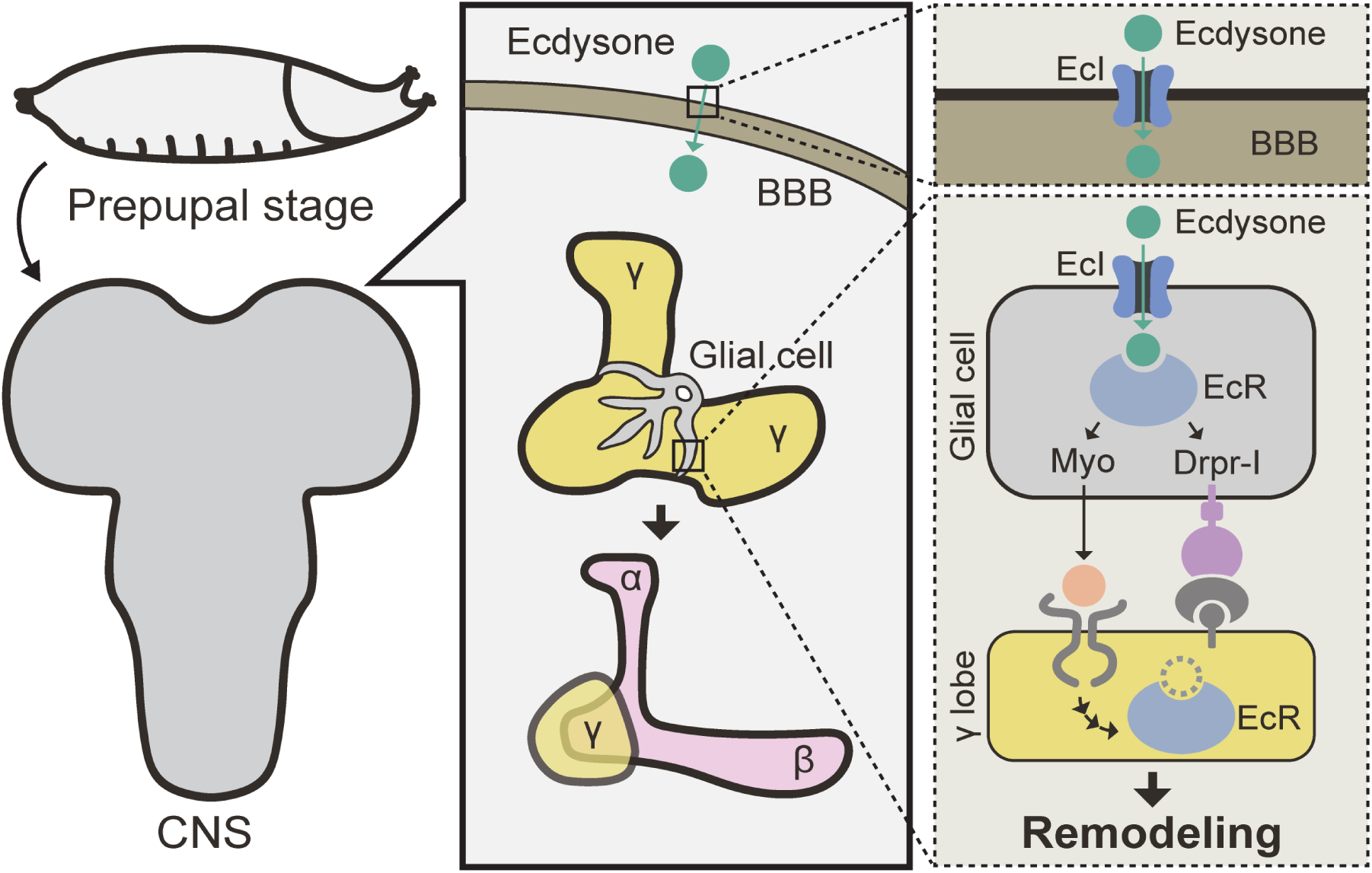
Schematic representation of the results obtained in this study. Ecdysone enters glial cells through the BBB via EcI and binds to EcR to induce expression of *myoglianin* (*myo*) and *draper isoform-I* (*drpr-I*). Drpr-I binds to its ligand presented by the MB γ axons to form a bridge and promotes glial engulfment. Myo is received by the MB and induces expression of EcR, which further facilitates MB remodeling potentially in a ligand-independent manner.

### EcI-independent EcR functions during CNS transformation

Dispensability of EcI and the other ecdysone importers (EcI-2, −3, and −4), but not EcR, in MB neurons (Figure 2) raises the possibility of cellular ecdysone uptake via alternative routes. However, the essential requirement of ecdysone importers in *Drosophila* and other insects (27) makes this scenario unlikely. *E23* overexpression, a manipulation that can effectively eliminate ecdysone action in other cellular populations within the CNS (Figure S2), has no discernible effects on MB remodeling (Figure 2E), making it further unlikely that a normal level of ecdysone is required in KCs for their remodeling. It is important to note here that there may still be a small, trace amount of ecdysone in KCs with *EcI* knockdown or *E23* overexpression, and such low-level ecdysone is critical for the EcR function in the MB during metamorphosis. Although such possibilities cannot be fully ruled out, potential ligand-independent functions of EcR need to be considered as an alternative possibility. Indeed, it has been previously shown that canonical ecdysone-inducible genes are not required for neuronal remodeling in the MB (14), consistent with their low expression in the CNS rescued by *myo* and *drpr-I* overexpression in glial cells (Figure 4E). As it is known that EcR acts as a transcription repressor in the absence of the ligand ecdysone (3, 44–46), it is possible that EcR is required in neurons to repress expression of genes that prevent neuronal remodeling. Alternatively, EcR can activate other signaling pathways in a ligand-independent manner (47), which in turn promotes neuronal remodeling. Although factors acting downstream of EcR in MB remodeling have been reported (9, 48), requirement of ecdysone for the induction of their expression has not been investigated. Along with more detailed investigations on these known EcR targets, transcriptome analyses of neurons undergoing remodeling from precisely staged samples, as was conducted in the *Drosophila* optic lobes at single-cell levels (20, 49), may provide more insight into molecular mechanisms of potential ligand-independent EcR functions in future studies.

Although glial overexpression of *myo* and *drpr* can dramatically rescue the CNS morphology defects and developmental lethality caused by blocking ecdysone entry into the CNS, our results indicate that ecdysone is also required in non-glial cells, such as the neuroepithelial cells in the optic lobes. Knocking down *EcI* in entire neurons using *nSyb-GAL4* also caused pharate adult lethality (Figure S1B), suggesting that ecdysone is required at least in some neurons to complete metamorphosis. Further investigation into the neurons that require ecdysone during metamorphosis is clearly warranted, which is expected to advance our understanding of how insect metamorphosis is regulated by interactions between the endocrine and nervous systems.

### Significance of ecdysone-myoglianin interaction during insect metamorphosis

Our *ex vivo* CNS culture experiments showed that *myo* is expressed in *Drosophila* glial cells in an ecdysone-dependent manner, suggesting that this TGF-β signaling pathway is acting downstream of ecdysone signaling. On the other hand, glia-derived myo upregulates *EcR* expression in MB neurons, although this EcR may act in a ligand-independent manner as discussed above. This bidirectional interaction between ecdysone and myo signaling pathways appears to be at the core of ecdysone-mediated CNS transformation during *Drosophila* metamorphosis. In this context, it is also noteworthy that myo has been shown to regulate biosynthesis of ecdysone as well as juvenile hormone, another important hormonal regulator of insect metamorphosis, in both hemimetabolous and holometabolous insects (50–52). Investigation of these multiple layers of ecdysone-myoglianin interactions in different orders of insects may help us reveal as-yet-unknown aspects of the hormonal regulation of insect metamorphosis, potentially from evolutionary perspectives.

### Hormonal gatekeeping at the BBB and brain development

Glial cells support mammalian brain development in multiple ways, and many of such glial cell functions, including the release of neurotrophic factors, regulation of neuroinflammation, and myelination, are regulated by hormones including steroids (53, 54). Importantly, thyroid hormones, mammalian lipophilic hormones that regulate oligodendrocyte differentiation and myelination, require their membrane transporters to be expressed in the BBB, whose deficiency causes neurological disorders (55–60). As illustrated by this example, hormonal gatekeeping at the BBB affects glial regulation of brain development in a wide variety of metazoans.

In insects, not only ecdysone but juvenile hormone entry into the CNS is also known to be controlled at the BBB, which affects behaviors in social insects (61, 62). Further elucidation of the molecular mechanisms of such hormonal gatekeeping at the insect BBB, therefore, may provide critical insights into how we can potentially manipulate brain development and other hormone-mediated processes, such as neuroinflammation and behavior, by controlling hormone entry into the animal brain.

## Materials and Methods

### Fly husbandry and stocks

All flies were raised at 25°C under 12 h/12 h light/dark cycle, except for those carrying *Repo-LexA*, which were reared at 21°C after hatching due to lethality at 25°C even in heterozygotes. The animals were reared on standard fly food containing 6 g *Drosophila* agar type II (Genesee Scientific, #66-103), 100 g D-(+)-glucose (SIGMA, #G8270-25KG), 50 g inactive dry yeast (Genesee Scientific, #62-106), 70 g yellow corn meal (Genesee Scientific, #62-101), 6 mL propionic acid (SIGMA, #402907-500ML), and 10 mL Tegosept (Genesee Scientific, #20-258) in 1,025 mL of water.

Flies used are as follows: *w^1118^* (#5905; the control strain), *nSyb-GAL4* (#51941), *Repo-GAL4* (#7415), *OK107-GAL4* (#854), *Ogre-GAL4* (#49340), *EcRE-LacZ* (#4516 and #4517), *UAS-EcR-RNAi* (#9327), *UAS-EcR-DN* (#6872), *UAS-mCD8-GFP* (#5130), *Df(2L)Exel6033* (#7516; deficiency allele over *EcI2*), *Df(2R)Exel7171* (#7902; deficiency allele over *EcI3* and *EcI4*), *R22B09-lexA* (#52600), *LexAop-mCD8-GFP* (32203), *UAS-drpr-I* (#67035), *UAS-EcR-B1* (#6469) and *Repo-LexA* (#604850) were obtained from the Bloomington *Drosophila* Stock Center (BDSC); *UAS-dicer2* (#60008 and #60009) was obtained from the Vienna *Drosophila* Resource Center (VDRC); *UAS-EcI-RNAi* (#7571R-1) was obtained from the National Institute of Genetics Fly Stock Center; *Elav-GAL4* and *UAS-myo^7D3^* were obtained from Michael B. O’Connor (University of Minnesota) (63). Ecdysone importer mutant alleles (*EcI2^1^*, *EcI3^1^*, and *EcI4^1^*) were generated previously (27). *UAS-E23*, *9-137-LexA*, *LexAop-EcI-RNAi,* and *LexAop- drpr-I* lines were generated in this study.

### Generation of transgenic lines

To generate *UAS-E23* line, *pUAST-E23* was prepared from the cDNA clone RE53253 from *Drosophila* Genomics Resource Center. A nonsense mutation in the clone (bp 2766) was corrected by site-directed mutagenesis as reported previously (64), and an *Xba*I site was introduced downstream of the start codon and used to insert a triple HA epitope. The product was then cloned into the *pUAST* vector, and transgenic flies were generated by BestGene Inc. *9-137-LexA* strain was generated by replacing the GAL4 sequence of the *9-137-GAL4* strain obtained from Ulrike Heberlein (Janelia Research Campus) with the LexA sequence using HACK method (65). The transformant was established by WellGenetics Inc. *LexAop-EcI-RNAi* was generated by inserting the double strand EcI sequence of *UAS-EcI-RNAi* (National Institute of Genetics: #7571R-1) into the *pUASTattB Drosophila* Gene Expression Vector and replacing the UAS sequence with the LexAop sequence. The vector was constructed by VectorBuilder Inc. (Vector ID: VB220329-1463vyp), and inserted into the attP40 or attP2 landing site by BestGene Inc. *LexAop-drpr-I* was generated by inserting the CDS of *drpr-I* into the *pUASTattB Drosophila* Gene Expression Vector and replacing the UAS sequence with the LexAop sequence. The vector was constructed by VectorBuilder Inc. (Vector ID: VB230903-1019mxf), and inserted into the attP40 or attP2 landing site by BestGene Inc.

### Immunostaining

The CNS was dissected in 1X phosphate-buffered saline (PBS) (Fisher BioReagents), fixed with 4% paraformaldehyde (Electron Microscopy Sciences) in PBS containing 0.1% Triton X-100 for 20 min at room temperature (RT), and washed multiple times with PBS containing 0.1% Triton X-100 (0.1% PBST). The tissues were blocked with 5% normal goat serum (NGS) (Sigma-Aldrich) in PBST for at least 1 h at RT, incubated overnight at 4°C with the primary antibody mix in PBST containing 5% NGS, washed multiple times with PBST, incubated for 2 h at RT with the secondary antibody mix in PBST containing 5% NGS, and washed multiple times again with PBST. When needed, DNA was stained with Hoechst 33342 (Thermo Fisher Scientific) at 1:2000 for 30 min at RT. After washing, tissues were mounted in Vectashield H-1000 (Vector Laboratories). Samples were visualized on a Zeiss Axio Imager M2 equipped with ApoTome.2 (Figure 2C, 2D, and S2B) or a Zeiss LSM 880 inverted confocal microscope with AiryScan (Figure 1B, 1C, 2B, 2E, 2F, S2C, S2E, 3F, 4D, S5A, and S6A).

Primary antibodies used were rat anti-N-cadherin (clone: DN-Ex #8, 1:50, Developmental Studies Hybridoma Bank), mouse anti-Fasciclin II (clone: 1D4, 1:500, Developmental Studies Hybridoma Bank), mouse anti-β-Galactosidase (LacZ) (clone: 40-1a, 1:500, Developmental Studies Hybridoma Bank), mouse anti-EcR (clone: Ag10.2, 1:200, Developmental Studies Hybridoma Bank), mouse anti-EcR-B1 (clone: AD4.4, 1:10, Developmental Studies Hybridoma Bank), mouse anti-GFP (clone: GFP-20, 1:1000, Sigma) and chicken anti-GFP (1:1000, Abcam). Secondary antibodies used were Alexa Fluor 488 goat anti-chicken (1:500, Thermo Fisher Scientific), Alexa Fluor 488 goat anti-mouse (1:500, Thermo Fisher Scientific), Alexa Fluor 488 goat anti-rat (1:500, Thermo Fisher Scientific), and Alexa Fluor 546 goat anti-mouse (1:500, Thermo Fisher Scientific).

### Scoring pupal lethality and eclosion rate

Egg laying was induced on 3% agar plates containing 30% grape-juice (Welch) at 25°C. Newly-hatched larvae were collected into food vials (100 larvae each) and raised on mashed fly food. Vials were incubated at 25°C for at least 14 days, after which dead pupae and emerged adults were counted.

### RNA extraction and quantitative reverse transcription PCR

The CNS was collected from 15 larvae or pupae in 1.5 ml tubes and flash-frozen immediately in liquid nitrogen. Total RNA of the collected CNS samples was extracted using TRIzol reagent (Invitrogen) according to the manufacturer’s instructions. Extracted RNA was further purified by RNeasy mini kit (Qiagen) following the manufacturer’s instructions, combined with treatment with RNase-Free DNase Set (Qiagen). cDNA was generated from purified total RNA using PrimeScript RT Master Mix (Takara Bio). qRT-PCR was performed on the CFX connect real-time PCR detection system (Bio-Rad) using SYBR Premix Ex Taq II (Tli RNaseH Plus) (Takara Bio). The amount of target RNA was normalized to *ribosomal protein 49 (rp49)* in the same samples, and relative fold changes were calculated. Three separate samples were collected for each experiment. The primers used to measure transcript levels are shown in Table S1. The primers to detect *E74A*, *E74B*, *E75B*, and *rp49* levels were previously reported (16).

### *Ex vivo* CNS culture with 20E

Mid-third instar larvae (24 hours after L3 ecdysis) were rinsed in PBS and dissected in Schneider’s *Drosophila* medium (GIBCO). The ring gland and imaginal disks attached to the CNS were removed. Dissected CNS was cultured in 3 mL of Schneider’s *Drosophila* medium containing 0.05% EtOH with 20E at various concentrations for 2 h in a sterile Petri dish (Olympus plastics) at 25°C.

### Image analysis

To quantify the size of the brain lobes or optic lobes in the CNS, the width of the lobe on each side was obtained from z stack images of the CNS using ZEN software (Carl Zeiss) and quantified using ImageJ software (NIH). Values were calculated relative to the mean of control groups. For the optic lobe size, the width of the optic lobe divided by the width of the brain lobe was used for the calculation.

### Statistical analyses

All statistical tests, significance levels, and sample sizes are reported in the figure legends. All experiments were performed at least 3 times independently. One-way or Two-way ANOVA with post-hoc Dunnett’s test was used for comparing multiple samples against a control. All statistical analyses were carried out in the “R” software environment (The R Foundation for Statistical Computing, Vienna, Austria). *p* ≤ 0.05 was considered significant. *P*-values are provided in comparison with control as ∗*p* ≤ 0.05, ∗∗*p* ≤ 0.01, ∗∗∗*p* ≤ 0.001, and “n.s.” for non-significant (p > 0.05).

## Data and code availability

This study did not generate any new computer code or algorithms. Raw images used for quantitative analyses are available from the corresponding author upon request.

## Acknowledgments

We are grateful to M.B. O’Connor (University of Minnesota, USA), J.H. Park (University of Tennessee, USA), the Bloomington *Drosophila* Stock Center (supported by NIH P40 OD018537), Vienna *Drosophila* Resource Center, National Institute of Genetics Fly Stock Center, *Drosophila* Genomics Resource Center (supported by NIH 2P40 OD010949), and Developmental Studies Hybridoma Bank for fly stocks and reagents, and FlyBase (supported by NIH U41 HG000739 and U24 HG010859) for providing curated *Drosophila* genome information. We also thank U. Heberlein (Janelia Research Campus) for helpful advice on the P element insertion site of *9-137-GAL4*. This study was supported by a Postdoctoral Fellowship for Research Abroad from the Japan Society for the Promotion of Science to N.O. and E.I., the Naito Foundation Subsidy for Dispatch of Young Researchers Abroad to N.O., the 10th Tomizawa Jun-ichi & Keiko Fund of MBSJ for Young Scientists to N.O., a Pew Biomedical Scholars Award from the Pew Charitable Trusts to N.Y., an NIH Director’s New Innovator Award DP2 GM132929 to N.Y., and an NIH grant R35 GM153331 from NIGMS to N.Y.

## Author Contributions

Conceptualization, E.I., N.O. and N.Y.; Methodology, E.I., N.O. and N.Y.; Investigation, E.I., N.O. and N.Y.; Writing – Review & Editing, E.I., N.O. and N.Y.; Supervision, N.Y.; Funding Acquisition, E.I., N.O. and N.Y.

## References

1. J. W. Truman, L. M. Riddiford, Endocrine insights into the evolution of metamorphosis in insects. Annu Rev Entomol 47, 467–500 (2002).

2. N. Yamanaka, Ecdysteroid signalling in insects—From biosynthesis to gene expression regulation. Adv In Insect Phys 60, 1–36 (2021).

3. J. W. Truman, L. M. Riddiford, Drosophila postembryonic nervous system development: a model for the endocrine control of development. Genetics 223 (2023).

4. J. W. Truman, Hormonal Control of the Form and Function of the Nervous System. Comprehensive Molecular Insect Science 2-6, 135–163 (2005).

5. J. W. Truman, Developmental neuroethology of insect metamorphosis. J Neurobiol 23, 1404–1422 (1992).

6. J. W. Truman, Steroid receptors and nervous system metamorphosis in insects. Dev Neurosci 18, 87–101 (1996).

7. C. Consoulas, C. Duch, R. J. Bayline, R. B. Levine, Behavioral transformations during metamorphosis: remodeling of neural and motor systems. Brain Res Bull 53, 571–583 (2000).

8. L. Veverytsa, D. W. Allan, Subtype-specific neuronal remodeling during Drosophila metamorphosis. Fly (Austin) 7, 78–86 (2013).

9. A. Boulanger, J. M. Dura, Nuclear receptors and Drosophila neuronal remodeling. Biochim Biophys Acta 1849, 187–195 (2015).

10. S. P. Yaniv, O. Schuldiner, A fly’s view of neuronal remodeling. Wiley Interdiscip Rev Dev Biol 5, 618–635 (2016).

11. M. H. Syed, B. Mark, C. Q. Doe, Playing Well with Others: Extrinsic Cues Regulate Neural Progenitor Temporal Identity to Generate Neuronal Diversity. Trends Genet 33, 933–942 (2017).

12. M. N. Modi, Y. Shuai, G. C. Turner, The Drosophila Mushroom Body: From Architecture to Algorithm in a Learning Circuit. Annu Rev Neurosci 43, 465–484 (2020).

13. R. L. Davis, Learning and memory using Drosophila melanogaster: a focus on advances made in the fifth decade of research. Genetics 224 (2023).

14. T. Lee, S. Marticke, C. Sung, S. Robinow, L. Luo, Cell-autonomous requirement of the USP/EcR-B ecdysone receptor for mushroom body neuronal remodeling in Drosophila. Neuron 28, 807–818 (2000).

15. T. Awasaki, T. Lee, Orphan nuclear receptors control neuronal remodeling during fly metamorphosis. Nat Neurosci 14, 6–7 (2011).

16. N. Okamoto, et al., A Membrane Transporter Is Required for Steroid Hormone Uptake in Drosophila. Dev Cell 47, 294–305.e7 (2018).

17. N. Okamoto, N. Yamanaka, Steroid Hormone Entry into the Brain Requires a Membrane Transporter in Drosophila. Curr Biol 30, 359–366.e3 (2020).

18. D. T. Champlin, J. W. Truman, Ecdysteroid control of cell proliferation during optic lobe neurogenesis in the moth Manduca sexta. Development 125, 269–277 (1998).

19. E. Lanet, A. P. Gould, C. Maurange, Protection of neuronal diversity at the expense of neuronal numbers during nutrient restriction in the Drosophila visual system. Cell Rep 3, 587–594 (2013).

20. S. Jain, et al., A global timing mechanism regulates cell-type-specific wiring programmes. Nature 603, 112–118 (2022).

21. M. R. Koelle, et al., The Drosophila EcR gene encodes an ecdysone receptor, a new member of the steroid receptor superfamily. Cell 67, 59–77 (1991).

22. T. Lee, S. Marticke, C. Sung, S. Robinow, L. Luo, Cell-autonomous requirement of the USP/EcR-B ecdysone receptor for mushroom body neuronal remodeling in Drosophila. Neuron 28, 807–818 (2000).

23. G. Marchetti, G. Tavosanis, Steroid Hormone Ecdysone Signaling Specifies Mushroom Body Neuron Sequential Fate via Chinmo. Curr Biol 27, 3017–3024.e4 (2017).

24. T. Hock, T. Cottrill, J. Keegan, D. Garza, The E23 early gene of Drosophila encodes an ecdysone-inducible ATP-binding cassette transporter capable of repressing ecdysone-mediated gene activation. Proc Natl Acad Sci U S A 97, 9519–9524 (2000).

25. T. Q. Itoh, T. Tanimura, A. Matsumoto, Membrane-bound transporter controls the circadian transcription of clock genes in Drosophila. Genes Cells 16, 1159–1167 (2011).

26. A. A. Evdokimova, et al., Transcriptional induction by ecdysone in Drosophila salivary glands involves an increase in chromatin accessibility and acetylation. Nucleic Acids Res 53, 284 (2025).

27. L. V. Hun, et al., Essential functions of mosquito ecdysone importers in development and reproduction. Proc Natl Acad Sci U S A 119 (2022).

28. D. T. Champlin, J. W. Truman, Ecdysteroid control of cell proliferation during optic lobe neurogenesis in the moth Manduca sexta. Development 125, 269–277 (1998).

29. D. T. Champlin, J. W. Truman, Ecdysteroid coordinates optic lobe neurogenesis via a nitric oxide signaling pathway. Development 127, 3543–3551 (2000).

30. P. C. H. Lo, M. Frasch, Sequence and expression of myoglianin, a novel Drosophila gene of the TGF-β superfamily. Mech Dev 86, 171–175 (1999).

31. X. Zheng, et al., TGF-β signaling activates steroid hormone receptor expression during neuronal remodeling in the Drosophila brain. Cell 112, 303–315 (2003).

32. T. Awasaki, Y. Huang, M. B. O’Connor, T. Lee, Glia instruct developmental neuronal remodeling through TGF-β signaling. Nat Neurosci 14, 821–823 (2011).

33. A. M. Rossi, C. Desplan, Extrinsic activin signaling cooperates with an intrinsic temporal program to increase mushroom body neuronal diversity. Elife 9, 1–23 (2020).

34. Z. Wang, G. Lee, R. Vuong, J. H. Park, Two-factor specification of apoptosis: TGF-β signaling acts cooperatively with ecdysone signaling to induce cell-and stage-specific apoptosis of larval neurons during metamorphosis in Drosophila melanogaster. Apoptosis 24, 972–989 (2019).

35. Y. Li, P. Haynes, S. L. Zhang, Z. Yue, A. Sehgal, Ecdysone acts through cortex glia to regulate sleep in Drosophila. Elife 12 (2023).

36. M. R. Freeman, J. Delrow, J. Kim, E. Johnson, C. Q. Doe, Unwrapping glial biology: Gcm target genes regulating glial development, diversification, and function. Neuron 38, 567–580 (2003).

37. Y. Hakim, S. P. Yaniv, O. Schuldiner, Astrocytes play a key role in Drosophila mushroom body axon pruning. PLoS One 9 (2014).

38. T. Awasaki, et al., Essential role of the apoptotic cell engulfment genes draper and ced-6 in programmed axon pruning during Drosophila metamorphosis. Neuron 50, 855–867 (2006).

39. E. D. Hoopfer, et al., Wlds protection distinguishes axon degeneration following injury from naturally occurring developmental pruning. Neuron 50, 883–895 (2006).

40. A. Boulanger, et al., Axonal chemokine-like Orion induces astrocyte infiltration and engulfment during mushroom body neuronal remodeling. Nat Commun 12 (2021).

41. H. Ji, et al., The Drosophila chemokine-like Orion bridges phosphatidylserine and Draper in phagocytosis of neurons. Proc Natl Acad Sci U S A 120 (2023).

42. S. Lin, The making of the Drosophila mushroom body. Front Physiol 14, 1091248 (2023).

43. M. A. Logan, et al., Negative regulation of glial engulfment activity by Draper terminates glial responses to axon injury. Nat Neurosci 15, 722–730 (2012).

44. H. Morrow, C. K. Mirth, Timing Drosophila development through steroid hormone action. Curr Opin Genet Dev 84 (2024).

45. J. Wardwell-Ozgo, et al., An EcR probe reveals mechanisms of the ecdysone-mediated switch from repression-to-activation on target genes in the larval wing disc. bioRxiv 2022.04.07.487542 (2022). 10.1101/2022.04.07.487542.

46. G. Perez-Mockus, et al., The Drosophila ecdysone receptor promotes or suppresses proliferation according to ligand level. Dev Cell 58, 2128–2139.e4 (2023).

47. A. Mansilla, F. A. Martin, D. Martin, A. Ferrus, Ligand-independent requirements of steroid receptors EcR and USP for cell survival. Cell Death Differ 23, 405–416 (2016).

48. F. Yu, O. Schuldiner, Axon and dendrite pruning in Drosophila. Curr Opin Neurobiol 27, 192–198 (2014).

49. M. N. Özel, et al., Neuronal diversity and convergence in a visual system developmental atlas. Nature 589, 88–95 (2021).

50. Y. Ishimaru, et al., TGF-β signaling in insects regulates metamorphosis via juvenile hormone biosynthesis. Proc Natl Acad Sci U S A 113, 5634–5639 (2016).

51. O. Kamsoi, X. Belles, Myoglianin triggers the premetamorphosis stage in hemimetabolan insects. FASEB J 33, 3659–3669 (2019).

52. S. Chafino, R. Salvia, J. Cruz, D. Martín, X. Franch-Marro, TGFß/activin-dependent activation of Torso controls the timing of the metamorphic transition in the red flour beetle Tribolium castaneum. PLoS Genet 19 (2023).

53. I. Lago-Baldaia, V. M. Fernandes, S. D. Ackerman, More Than Mortar: Glia as Architects of Nervous System Development and Disease. Front Cell Dev Biol 8 (2020).

54. C. Guevara, F. Ortiz, Glial-derived transforming growth factor β1 (TGF-β1): a key factor in multiple sclerosis neuroinflammation. Neural Regen Res 16, 510–511 (2021).

55. W. E. Visser, E. C. H. Friesema, T. J. Visser, Minireview: thyroid hormone transporters: the knowns and the unknowns. Mol Endocrinol 25, 1–14 (2011).

56. J. Bernal, A. Guadaño-Ferraz, B. Morte, Thyroid hormone transporters-functions and clinical implications. Nat Rev Endocrinol 11, 406–417 (2015).

57. K. Landers, K. Richard, Traversing barriers - How thyroid hormones pass placental, blood-brain and blood-cerebrospinal fluid barriers. Mol Cell Endocrinol 458, 22–28 (2017).

58. P. Strømme, et al., Mutated Thyroid Hormone Transporter OATP1C1 Associates with Severe Brain Hypometabolism and Juvenile Neurodegeneration. Thyroid 28, 1406–1415 (2018).

59. J. Bernal, New insights on thyroid hormone and the brain. Curr Opin Endocr Metab Res 2, 24–28 (2018).

60. J. Bernal, Thyroid Hormones in Brain Development and Function. Endotext (2022).

61. B. M. Jones, et al., Convergent and complementary selection shaped gains and losses of eusociality in sweat bees. Nat Ecol Evol 7, 557–569 (2023).

62. L. Ju, et al., Hormonal gatekeeping via the blood-brain barrier governs caste-specific behavior in ants. Cell 186, 4289–4309.e23 (2023).

63. S. C. Gesualdi, T. E. Haerry, Distinct signaling of Drosophila Activin/TGF-beta family members. Fly (Austin) 1, 212–221 (2007).

64. N. Yamanaka, G. Marqués, M. B. O’Connor, Vesicle-Mediated Steroid Hormone Secretion in Drosophila melanogaster. Cell 163, 907–919 (2015).

65. C. C. Lin, C. J. Potter, Editing transgenic DNA components by inducible gene replacement in Drosophila melanogaster. Genetics 203, 1613–1628 (2016).

