## Supporting Information for "Steroid hormone-dependent glial-neuronal interaction promotes brain development during *Drosophila* metamorphosis"

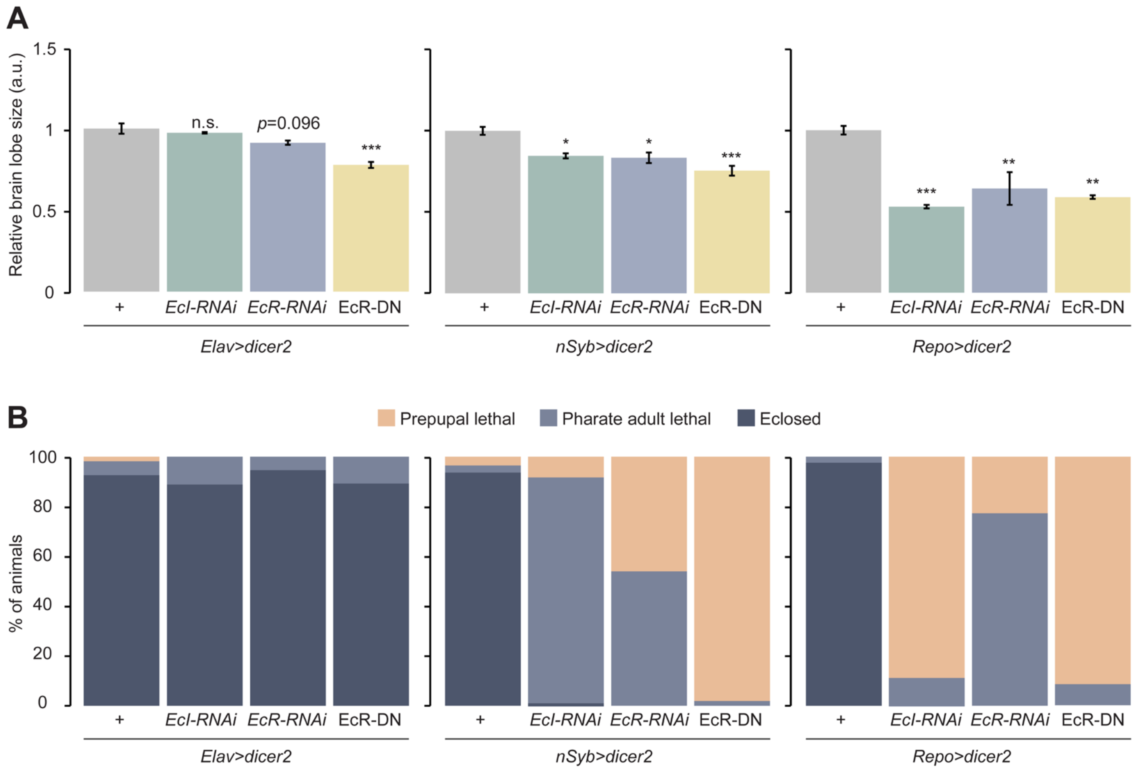


**Figure S1. Quantitative analyses of CNS transformation defects and developmental lethality induced by disrupting ecdysone signaling.**

(A, B) Relative brain lobe size (A) at 24 h after puparium formation (24 hAPF) and developmental lethality (B) of control animals, as well as those expressing *EcI-RNAi*, *EcR-RNAi*, or a dominant-negative form of *EcR* (*EcR-DN*) in neurons or glial cells. *Elav-GAL4, UAS-dicer2* or *nSyb-GAL4, UAS-dicer2* and *Repo-GAL4, UAS-dicer2* were used to induce transgene expression in entire neurons and glial cells, respectively.

Values in A are relative to that of the control animals. a.u., arbitrary unit. One-way ANOVA with post-hoc Dunnett's test was used for A. $*p \leq 0.05$, $**p \leq 0.01$, $***p \leq 0.001$, and n.s. for non-significant ($p > 0.05$). All values in A are the means ± standard error ($n \geq 4$). Orange, light blue, and dark blue columns in B represent animals that died as prepupae, died as pharate adults, and eclosed normally, respectively.

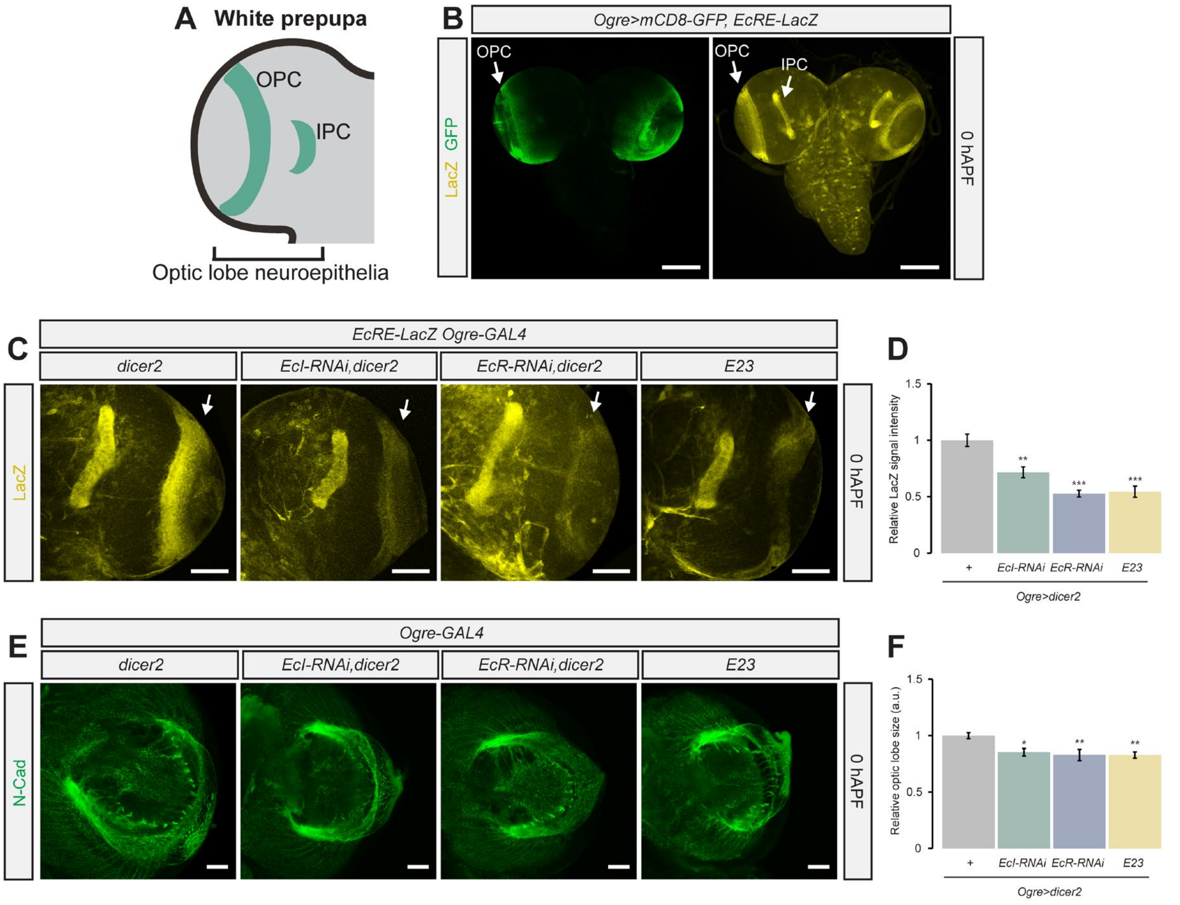


**Figure S2. Ecdysone is required in the optic lobes for cellular differentiation.**

(A) Schematic illustration of the optic lobe in the prepupal brain. OPC and IPC represent the outer and inner proliferation centers of the optic lobe neuroepithelia, respectively.

(B) Images of the prepupal brain at 0 h after puparium formation (0 hAPF). The OPCs were labeled with mCD8-GFP (green) driven by *Ogre-GAL4*, whereas ligand-dependent EcR activity was visualized by the ecdysone response element-driven LacZ reporter (*EcRE-LacZ*; yellow). Scale bars represent 100 µm.

(C) *EcRE-LacZ* reporter activity (yellow) in the prepupal brain at 0 hAPF. *Ogre-GAL4* was used to induce *EcI-RNAi*, *EcR-RNAi*, or *E23* overexpression in the OPCs. Scale bars represent 50 µm.

(D) Relative LacZ signal intensity at 0 hAPF of control animals, as well as those expressing *EcI-RNAi*, *EcR-RNAi*, or *E23* in the OPCs. Values are relative to that of the control animals.

(E) Images of the prepupal brain stained with anti-N-cadherin (N-Cad) antibody (green) at 0 hAPF. *Ogre-GAL4* was used to induce *EcI-RNAi*, *EcR-RNAi*, or *E23* overexpression in the OPCs. Scale bars represent 25 µm.

(F) Relative optic lobe size at 0 hAPF of control animals, as well as those expressing *EcI-RNAi*, *EcR-RNAi*, or *E23* in the OPCs. Values are relative to that of the control animals. a.u., arbitrary unit.

One-way ANOVA with post-hoc Dunnett's test was used for D and F. $*p \leq 0.05$, $**p \leq 0.01$, $***p \leq 0.001$, and n.s. for non-significant ($p > 0.05$). All values are the means ± standard error ($n \geq 4$).

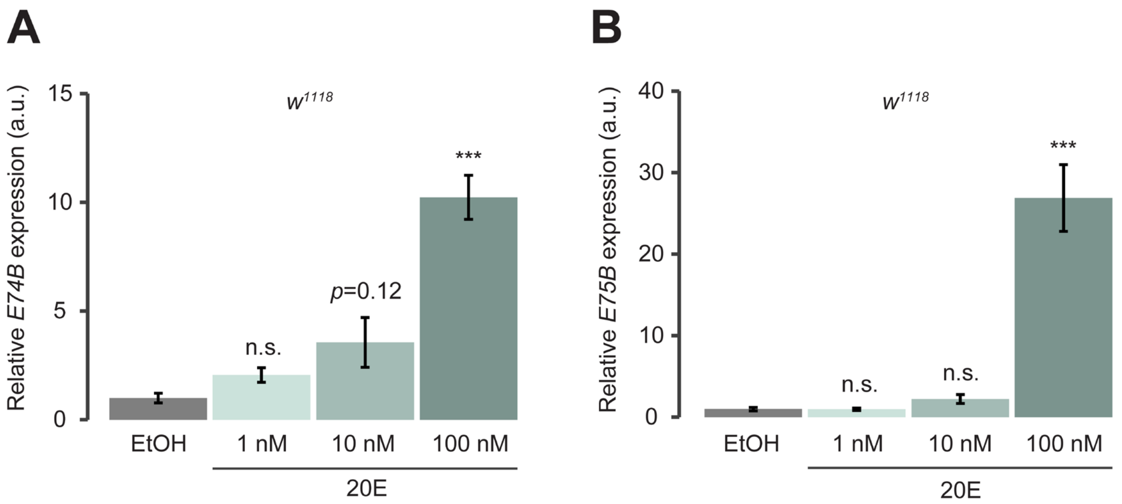


**Figure S3. Ecdysone-dependent induction of two ecdysone-responsive genes in the CNS cultured *ex vivo*.**

(A, B) Relative expression levels of *E74B* (A) and *E75B* (B) in the CNS of mid-L3 $w^{1118}$ larvae cultured *ex vivo* for 2 h with various concentrations of 20-hydroxyecdysone (20E). Values are relative to that of the solvent control (EtOH). a.u., arbitrary unit. One-way ANOVA with post-hoc Dunnett's test was used. $***p \leq 0.001$, and n.s. for non-significant ($p > 0.05$). All values are the means ± standard error ($n \geq 4$).

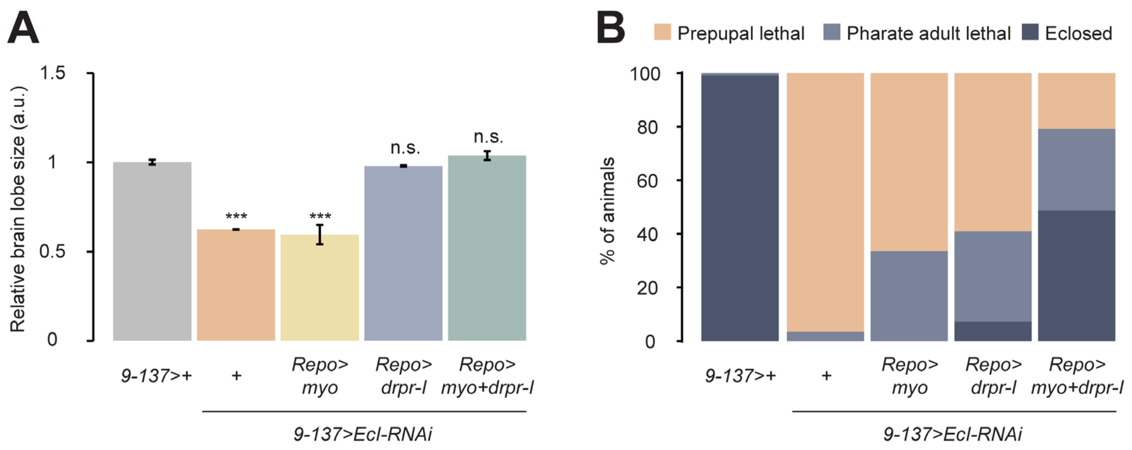


**Figure S4. Myoglianin- and Draper-mediated rescue of CNS transformation defects and developmental lethality induced by blocking ecdysone entry into the CNS.**

(A, B) Relative brain lobe size (A) at 24 h after puparium formation (24 hAPF) and developmental lethality (B) of control (*9-137*>+), as well as those expressing *EcI-RNAi* in the BBB with or without *myoglianin* (*myo*), *draper isoform-I* (*drpr-I*), or *myo*+*drpr-I* overexpression in glial cells.

Values in A are relative to that of *9-137*>+. a.u., arbitrary unit. One-way ANOVA with post-hoc Dunnett's test was used for A. ∗∗∗$p \leq 0.001$, and n.s. for non-significant ($p > 0.05$). All values in A are the means ± standard error ($n \geq 4$). Orange, light blue, and dark blue columns in B represent animals that died as prepupae, died as pharate adults, and eclosed normally, respectively.

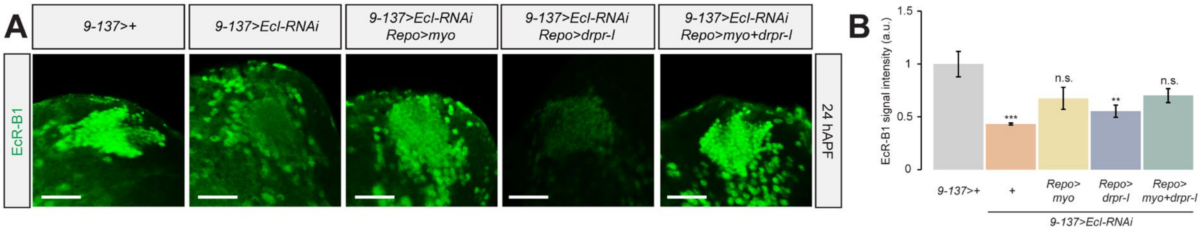


**Figure S5. Myoglianin-mediated rescue of EcR-B1 expression in Kenyon Cells.**

(A) Images of Kenyon cells stained with anti-EcR-B1 antibody (green) at 0 h after puparium formation (0 hAPF). *9-137-LexA* was used to induce *Ecl-RNAi* in the BBB, whereas *Repo-GAL4* was used to overexpress *myoglianin* (*myo*), *draper isoform-I* (*drpr-I*), or *myo+drpr-I* in glial cells. Scale bars represent 25 μm.

(B) Relative signal intensity of anti-EcR-B1 at 0 hAPF of control (*9-137*>+), as well as those expressing *Ecl-RNAi* in the BBB with or without *myo*, *drpr-I*, or *myo+drpr-I* overexpression in glial cells. Values are relative to that of *9-137*>+. a.u., arbitrary unit. One-way ANOVA with post-hoc Dunnett's test was used. $**p \leq 0.01$, $***p \leq 0.001$, and n.s. for non-significant ($p > 0.05$). All values are the means ± standard error ($n \geq 4$).

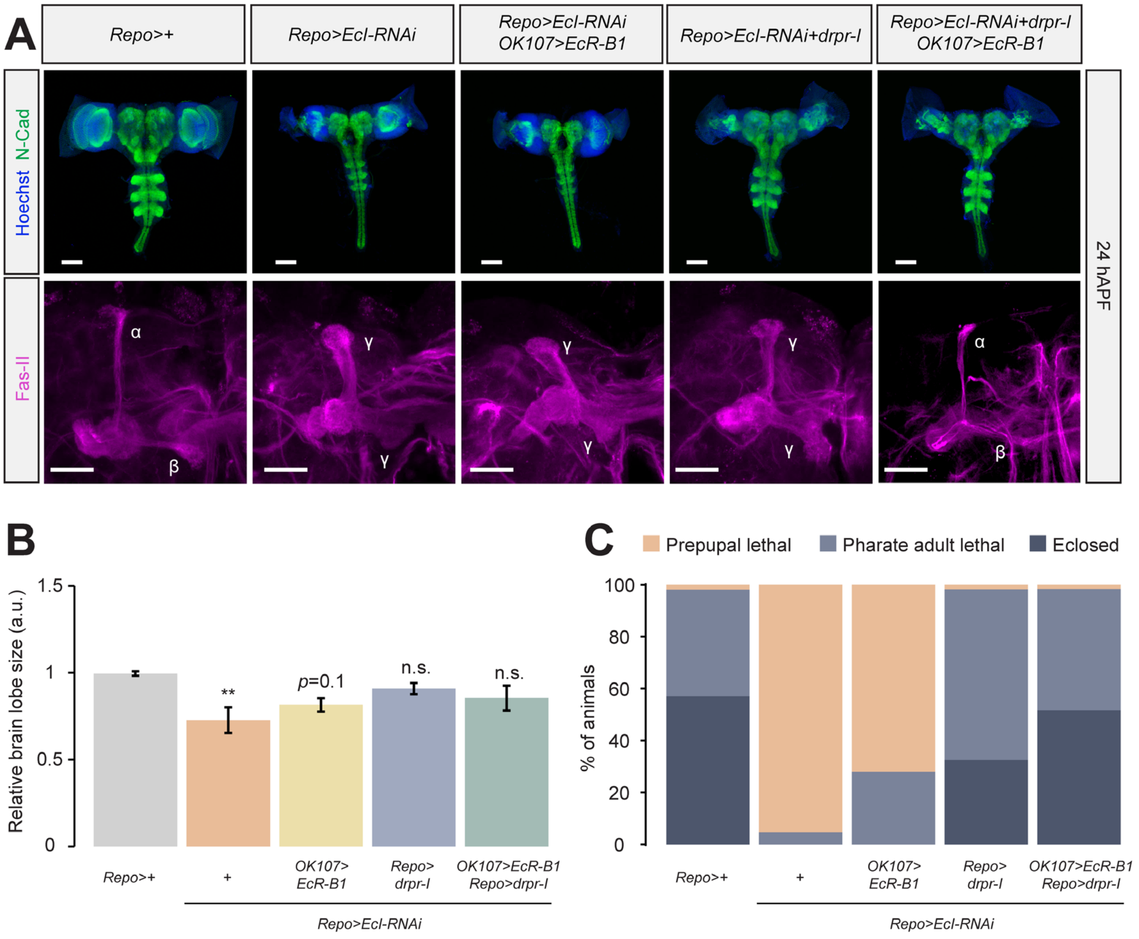


**Figure S6. EcR-B1- and Draper-mediated rescue of CNS transformation defects and developmental lethality induced by blocking ecdysone entry into the CNS.**

(A) Images of the pupal CNS and mushroom body (MB) at 24 h after puparium formation (24 hAPF). *Repo-LexA* was used to induce *EcI-RNAi* and to overexpress *draper isoform-I* (*drpr-I*) in the glial cells, whereas *OK107-GAL4* was used to overexpress *EcR-B1* in the MB. Neuronal axons were stained with anti-N-cadherin (N-Cad) antibody (green), nuclei were stained with Hoechst 33342 (blue), and the MB lobes were visualized by anti-Fasciclin-II (Fas-II) antibody staining (magenta). Scale bars represent 100 μm for top and 25 μm for bottom panels.

(B, C) Relative brain lobe size (B) at 24 hAPF and developmental lethality (C) of control (*Repo*>+), as well as those expressing *EcI-RNAi* in the glial cells with or without *EcR-B1* and/or *drpr-I* overexpression in the MB and in glial cells, respectively.

Values in B are relative to that of *Repo*>+. a.u., arbitrary unit. One-way ANOVA with post-hoc Dunnett's test was used for B. $**p \leq 0.01$, and n.s. for non-significant ($p > 0.05$). All values in B are the means ± standard error ($n \geq 4$). Orange, light blue, and dark blue columns in C represent animals that died as prepupae, died as pharate adults, and eclosed normally, respectively.

**Table 1. Primers used in this study.**

| Primers for qRT-PCR | | | |
|---|---|---|---|
| **Gene Name** | **CG Number** | **Forward (5'-3')** | **Reverse(5'-3')** |
| myo (Common) | CG1838 | ACCAACGATGAGGAGTACGAG | ACCTTGCGATTGTGCCGAAC |
| drpr (Isoform-I) | CG2086 | ACTTCAACAACAGCCTGGCC | AGTAACCGCTCCGACAGTTG |
